# Predicting *Pseudomonas aeruginosa* drug resistance using artificial intelligence and clinical MALDI-TOF mass spectra

**DOI:** 10.1101/2023.10.25.563934

**Authors:** Hoai-An Nguyen, Anton Y. Peleg, Jiangning Song, Bhavna Antony, Geoffrey I. Webb, Jessica A. Wisniewski, Luke V. Blakeway, Gnei Z. Badoordeen, Ravali Theegala, Helen Zisis, David L. Dowe, Nenad Macesic

**Affiliations:** Department of Infectious Diseases, The Alfred Hospital and Central Clinical School, Monash University, Melbourne, Australia; Monash Biomedicine Discovery Institute, Department of Microbiology, Monash University, Melbourne, Australia; Centre to Impact AMR, Monash University, Melbourne, Australia; Monash Biomedicine Discovery Institute, Department of Biochemistry & Molecular Biology, Monash University, Melbourne, Australia; Department of Data Science & AI, Monash University, Melbourne, Australia

**Keywords:** antimicrobial resistance, MALDI-TOF MS, machine learning, *Pseudomonas aeruginosa*

## Abstract

Matrix-assisted laser desorption/ionization–time of flight mass spectrometry (MALDI-TOF MS) is widely used in clinical microbiology laboratories for bacterial identification but its use for prediction of antimicrobial resistance (AMR) remains limited. Here, we used MALDI-TOF MS with artificial intelligence (AI) approaches to successfully predict AMR in *Pseudomonas aeruginosa*, a priority pathogen with complex AMR mechanisms. The highest performance was achieved for modern β-lactam/β-lactamase inhibitor drugs, namely ceftazidime/avibactam and ceftolozane/tazobactam, with area under the receiver operating characteristic curve (AUROC) of 0.86 and 0.87, respectively. As part of this work, we developed dynamic binning, a feature engineering technique that effectively reduces the high-dimensional feature set and has wide-ranging applicability to MALDI-TOF MS data. Compared to conventional methods, our approach yielded superior performance in 10 of 11 antimicrobials. Moreover, we showcase the efficacy of transfer learning in enhancing the performance for 7 of 11 antimicrobials. By assessing the contribution of features to the model’s prediction, we identified proteins that may contribute to AMR mechanisms. Our findings demonstrate the potential of combining AI with MALDI-TOF MS as a rapid AMR diagnostic tool for *Pseudomonas aeruginosa*.

## Introduction

*Pseudomonas aeruginosa* is an opportunistic pathogen that causes significant global morbidity and mortality [1] and has been identified by the World Health Organization (WHO) as a critical priority pathogen [2]. The development of antimicrobial resistance (AMR) in *P. aeruginosa* is often due to a complex interplay of intrinsic mechanisms, chromosomal mutations, and the ability to horizontally acquire resistance determinants from other species [3–5]. This diverse repertoire can lead to rapid development of AMR to different antimicrobial agents, including last resort treatments for *P. aeruginosa* infections. Furthermore, the global spread of multidrug- resistant, high-risk clones such as ST235, ST111, or ST233 has made treating *P. aeruginosa* increasingly challenging [6]. The lack of new antimicrobial discoveries makes optimizing use of current antipseudomonal agents an urgent priority.

Treatment of *P. aeruginosa* infection often begins empirically with later adjustment based on results of antimicrobial susceptibility testing (AST). Accurate and rapid AST methods are thus critical in selecting appropriate antimicrobial agents, both to effectively treat potentially life- threatening infections and to reduce the risk of inducing AMR due to inappropriate antimicrobial exposure. Traditional culture-based AST methods are accurate, but their turnaround time can often be more than 72-96 hours, leading to unacceptable delays in treatment [7].

In the face of these challenges, matrix-assisted laser desorption/ionization-time of flight mass spectrometry (MALDI-TOF MS) has emerged as a key technology in clinical microbiology. The technique characterizes the protein profile of a particular pathogen based on the spectrum recorded mass and quantity of ionized particles. MALDI-TOF MS has already been implemented in clinical microbiology laboratories for routine bacterial identification [8, 9] and investigation is ongoing for its potential use for AST by comparing the spectra generated with known AMR patterns [10]. Current approaches for use of MALDI-TOF MS in this setting rely on matches to a database, limiting their applicability for complex AMR mechanisms resulting from multiple interacting factors. Recent advancements in machine learning (ML) algorithms and computing resources make artificial intelligence (AI) approaches a promising way to predict AMR using MALDI-TOF MS data. While this has been utilized in other key bacterial pathogens such as methicillin resistant *Staphylococcus aureus* and vancomycin resistant *Enterococcus* species, the technique has yet to be investigated in *P. aeruginosa* [11–15].

To address this gap, we applied ML approaches to predict AMR in *P. aeruginosa* with broader applicability to other use cases relying on analysis of MALDI-TOF spectra. We present two methods aimed at enhancing the predictive performance. Firstly, we developed dynamic binning, a novel and simple feature engineering technique, which yielded superior results compared to conventional approaches. Moreover, through the analysis of salient feature bins, we identified potential AMR markers for *β*-lactam antimicrobials. Finally, we used transfer learning to acquire an embedding layer from a large publicly available MALDI-TOF MS dataset and effectively integrate these features into our model, resulting in improved performance.

## Results

### Data collection and resistance profiles

We generated MALDI-TOF spectra (Bruker Daltonics) on 380 *P. aeruginosa* isolates from our institution collected from 2004-2021. Following this, we conducted phenotypic AST using broth microdilution. The resulting minimum inhibitory concentrations (MICs) for 11 anti-pseudomonal antimicrobials were categorized in binary labels as ‘resistant’ and ‘nonresistant’ (Fig. 1) according to established breakpoints [16].

**Fig. 1.**
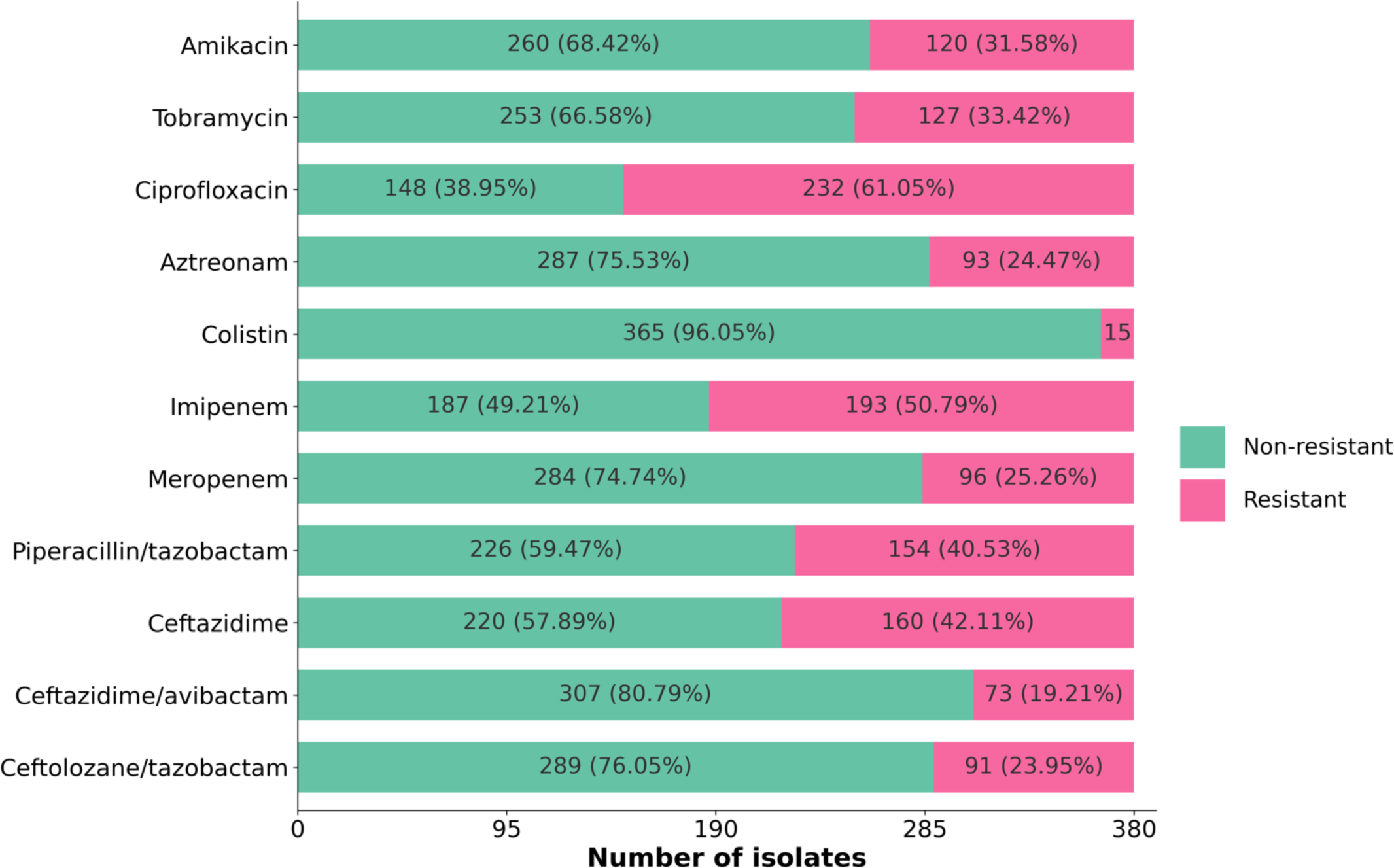
- Summary of antimicrobial susceptibility testing phenotypes. Number and proportion of non-resistant and resistant isolates of each antimicrobial. Broth microdilution assays were conducted for all 380 isolates to obtain MIC values for 11 antipseudomonal agents.

### Developing machine learning models for MALDI-TOF MS-based AMR prediction

We developed ML models using MALDI-TOF spectra as inputs to predict phenotypic susceptibility categories for each isolate and antimicrobial combination (Fig. 2). Spectra were initially preprocessed using a previously described approach [17]. Next, we introduced a novel, yet simple method called dynamic binning to extract features from the preprocessed spectra. Dynamic binning addresses the shortcomings of two commonly used feature engineering methods for MALDI-TOF MS data: peak detection (i.e., using intensity peaks as features) [13, 15, 18], and fixed-length binning (i.e., using bins of pre-specified size as features) [14, 17]. The former approach requires additional analysis steps for peak alignment and potentially overlooks regions without peaks, while the latter approach often generates a large feature set that greatly outnumbers sample size, thereby increasing the chances of over-fitting. With dynamic binning, we consider the quantity of intensity peaks in each region to dynamically determine the bin width for the binning process (see Methods). This allowed us to generate an 828-dimensional representation vector for each isolate.

**Fig. 2.**
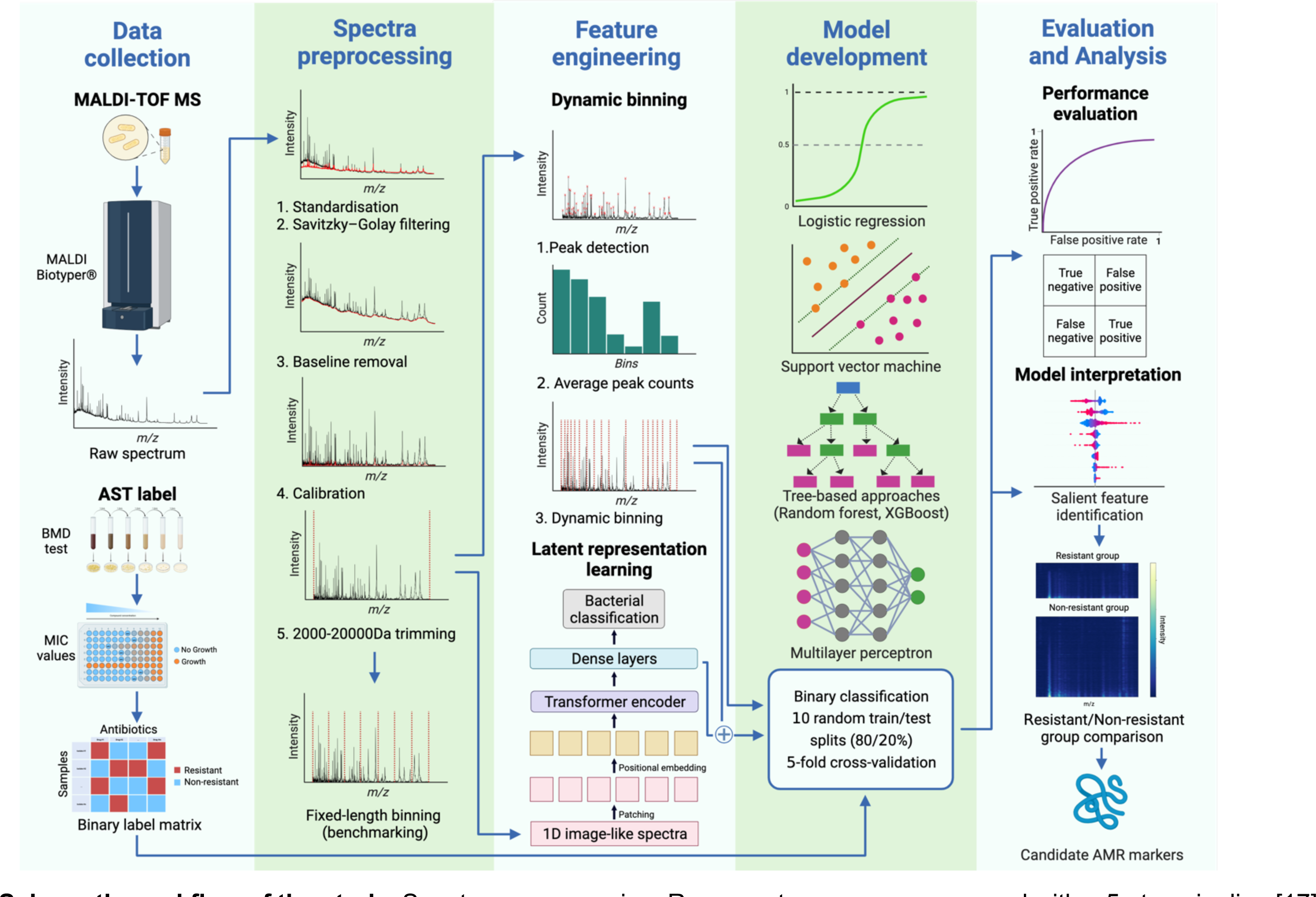
- Schematic workflow of the study. Spectra preprocessing: Raw spectra were preprocessed with a 5-step pipeline [17]. To generate the benchmarking feature sets, we conducted fixed-length binning (3-Da and 20-Da) of preprocessed spectra, resulting in feature vectors of 6,000 and 900 dimensions, respectively. Dynamic binning: we determined the mean number of intensity peaks in 500-Da intervals from 2,000 to 20,000 Da. Bin widths used for each region are inversely proportional to the corresponding peak count. Latent representation learning: a vision transformer model was trained to differentiate high-risk bacterial pathogens [19]. The final hidden layer of the model was used as embedding to extract additional features from the MALDI-TOF spectra. Model development: We used 5-fold cross validation with 80%-20% training-testing split to train models. Performance evaluation: AUROC was the key performance metric with log-loss, F1 score, precision, and recall also reported. Model interpretation: The Shapley Additive Explanations (SHAP) algorithm was used to identify feature bins with high importance [20]. We then analyzed the regions of interest to interpret models and identify potential biological AMR determinants. Abbreviations: MALDI-TOF MS - matrix-assisted laser desorption/ionization-time of flight mass spectrometry; AST – antimicrobial susceptibility testing; BMD – broth microdilution; MIC – minimum inhibitory concentration; m/z – mass-to-charge ratio.

For the prediction task, we employed five distinct supervised machine learning approaches (logistic regression [LR], random forest [RF], support vector machine [SVM], XGBoost [XGB], and multilayer perceptron [MLP]) and evaluated several ML performance metrics (see Methods). We trained and tested our models with 10 random splits and calculated the average metric values of all runs for each model and each antimicrobial.

### Successful prediction of antimicrobial susceptibility in *P. aeruginosa* using MALDI-TOF MS

The best predictive performance for each antimicrobial is presented in Fig. 3. Notably, our models performed best for novel *β*-lactam/*β*-lactamase inhibitor agents, achieving area under the receiver operating characteristic curve (AUROC) (mean, 95% CI) of 0.86 (0.74-0.98) and 0.87 (0.79-0.95) for ceftazidime/avibactam and ceftolozane/tazobactam, respectively. Performance remained high for aminoglycosides (amikacin and tobramycin) and ciprofloxacin, with AUROC of 0.84 (0.78-0.9), 0.82 (0.70-0.94), and 0.82 (0.72-0.92), respectively. Our results for other *β*-lactams were mixed, ranging from 0.69 (0.59-0.79) to 0.79 (0.65-0.93). For colistin, our models naively classified most isolates as non-resistant to colistin likely due to the highly imbalanced dataset (only 15/380 resistant isolates). SVM and RF were the algorithms that most frequently achieved highest AUROC (4/11 antimicrobials each).

**Fig. 3.**
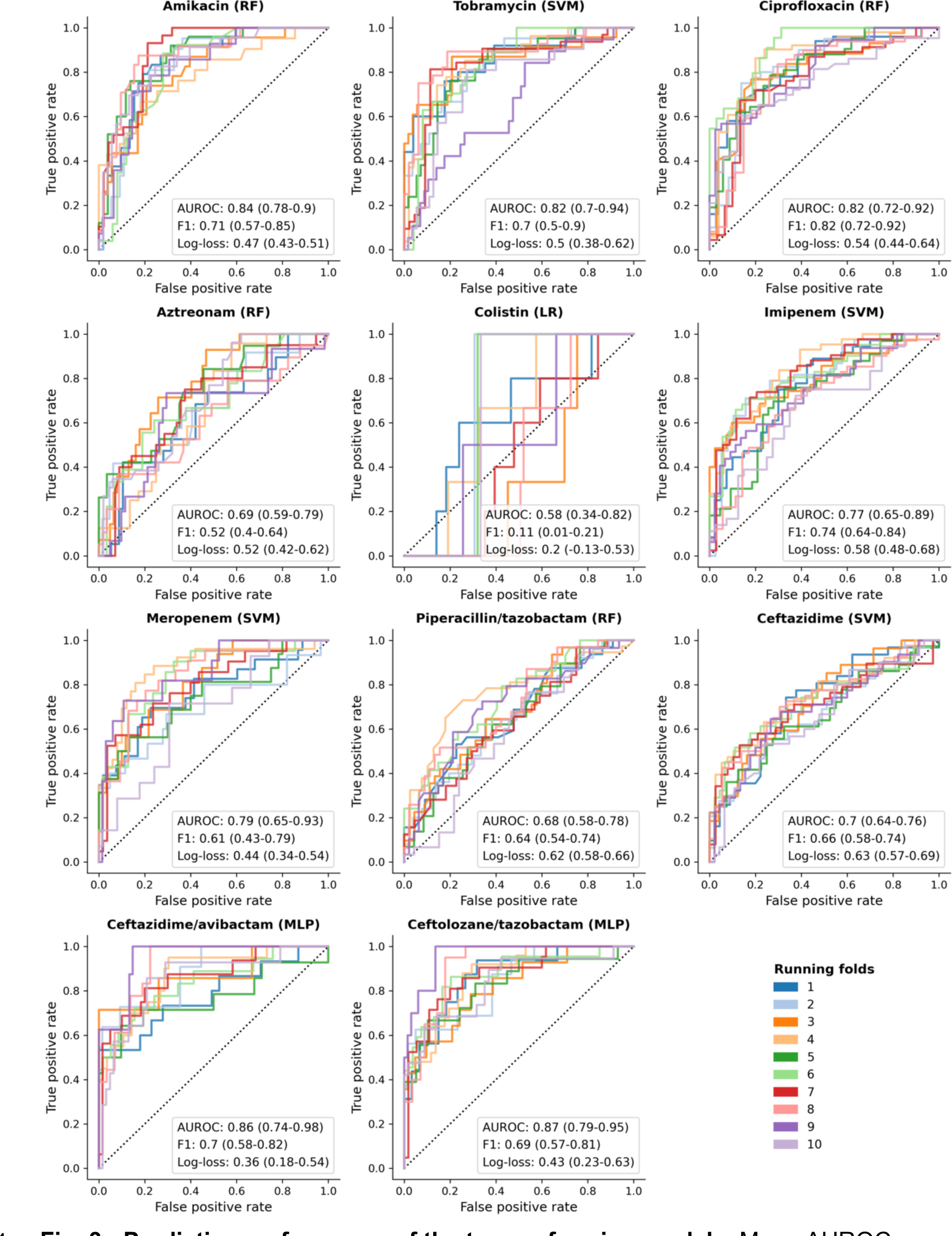
- Predictive performance of the top-performing models. Mean AUROC scores across 10 random splits are used to determine the best performing models. For each antimicrobial, the best performing algorithm is shown in parentheses after the antimicrobial. The plot represents the receiver-operating characteristic curves of the best model. Additionally, the corresponding F1 scores and log-loss values are included in the inset. All metrics are reported with mean values and 95% confidence intervals. Supplementary File 1 provides detailed metric values of the best performance models. Abbreviations: AUROC – area under the receiver-operating characteristic curve; LR: logistic regression; MLP: multilayer perceptron; RF: random forest; SVM: support vector machine.

### Dynamic binning method allows dimensionality reduction without reducing performance

We conducted a comparative analysis of dynamic binning with two fixed-length binning approaches: 3-Da binning (yielding 6000 features) [17], and 20-Da binning (yielding 900 features – a feature vector dimension close to that of dynamic binning). For comparison purposes, results for colistin were excluded because all methods performed poorly. Dynamic binning produced equal or superior results compared to both fixed-length binning settings for all 10 antimicrobials evaluated, despite using the smallest feature set (Table 1). In turn, 3-Da binning generally performed better (8/10 antimicrobials) or equally well (2/10 antimicrobials) to 20-Da binning, indicating that a simple reduction technique does not necessarily improve model performance.

**Table 1.** - Predictive performance of dynamic binning in comparison with fixed length binning. For each binning method and each antimicrobial, we report the best performing model and the corresponding AUROC, F1 score, and log-loss with 95% confidence intervals (CI). Bold text indicates models achieving the highest AUROC performance for each antimicrobial. Supplemental File 1 provides detailed performance of all models and metrics. Abbreviations: AUROC – area under the receiver-operating characteristic curve; LR: logistic regression; MLP: multilayer perceptron; RF: random forest; SVM: support vector machine.

| Antimicrobial | Dynamic binning |  |  |  | 3-Da binning |  |  |  | 20-Da binning |  |  |  |
| --- | --- | --- | --- | --- | --- | --- | --- | --- | --- | --- | --- | --- |
|  | Model | AUROC | F1 | Log-loss | Model | AUROC | F1 | Log-loss | Model | AUROC | F1 | Log-loss |
| Amikacin | **RF** | **0.84**<br>(0.78-0.9) | **0.71**<br>(0.57-0.85) | **0.47**<br>(0.43-0.51) | SVM | 0.82<br>(0.76-0.88) | 0.69<br>(0.57-0.81) | 0.48<br>(0.42-0.54) | RF | 0.83<br>(0.75-0.91) | 0.69<br>(0.57-0.81) | 0.48<br>(0.42-0.54) |
| Tobramycin | **SVM** | **0.82**<br>(0.7-0.94) | **0.7**<br>(0.5-0.9) | **0.5**<br>(0.38-0.62) | RF | 0.81<br>(0.71-0.91) | 0.66<br>(0.52-0.8) | 0.49<br>(0.41-0.57) | SVM | 0.8<br>(0.74-0.86) | 0.67<br>(0.55-0.79) | 0.5<br>(0.42-0.58) |
| Ciprofloxacin | **RF** | **0.82**<br>(0.72-0.92) | **0.82**<br>(0.72-0.92) | **0.54**<br>(0.44-0.64) | RF | 0.81<br>(0.71-0.91) | 0.82<br>(0.72-0.92) | 0.54<br>(0.44-0.64) | RF | 0.8<br>(0.7-0.9) | 0.82<br>(0.72-0.92) | 0.55<br>(0.45-0.65) |
| Aztreonam | **RF** | **0.69**<br>(0.59-0.79) | **0.52**<br>(0.4-0.64) | **0.52**<br>(0.42-0.62) | **RF** | **0.69**<br>(0.59-0.79) | **0.51**<br>(0.41-0.61) | **0.52**<br>(0.4-0.64) | RF | 0.67<br>(0.55-0.79) | 0.51<br>(0.39-0.63) | 0.56<br>(0.38-0.74) |
| Colistin | LR | 0.58<br>(0.34-0.82) | 0.11<br>(0.01-0.21) | 0.2<br>(-0.13-0.53) | **LR** | **0.63**<br>(0.28-0.98) | **0.14**<br>(0.04-0.24) | **0.25**<br>(-0.18-0.68) | LR | 0.59<br>(0.32-0.86) | 0.13<br>(-0.01-0.27) | 0.3<br>(-0.23-0.83) |
| Imipenem | **SVM** | **0.77**<br>(0.65-0.89) | **0.74**<br>(0.64-0.84) | **0.58**<br>(0.48-0.68) | **SVM** | **0.77**<br>(0.67-0.87) | **0.75**<br>(0.65-0.85) | **0.59**<br>(0.51-0.67) | SVM | 0.76<br>(0.66-0.86) | 0.74<br>(0.64-0.84) | 0.59<br>(0.49-0.69) |
| Meropenem | **SVM** | **0.79**<br>(0.65-0.93) | **0.61**<br>(0.43-0.79) | **0.44**<br>(0.34-0.54) | **RF** | **0.79**<br>(0.67-0.91) | **0.62**<br>(0.44-0.8) | **0.43**<br>(0.31-0.55) | LR | 0.77<br>(0.61-0.93) | 0.6<br>(0.44-0.76) | 0.59<br>(0.53-0.65) |
| Piperacillin/tazobactam | **RF** | **0.68**<br>(0.58-0.78) | **0.64**<br>(0.54-0.74) | **0.62**<br>(0.58-0.66) | SVM | 0.66<br>(0.54-0.78) | 0.64<br>(0.54-0.74) | 0.65<br>(0.61-0.69) | **SVM** | **0.68**<br>(0.58-0.78) | **0.65**<br>(0.57-0.73) | **0.64**<br>(0.58-0.7) |
| Ceftazidime | **SVM** | **0.7**<br>(0.64-0.76) | **0.66**<br>(0.58-0.74) | **0.63**<br>(0.57-0.69) | LR | 0.69<br>(0.63-0.75) | 0.66<br>(0.58-0.74) | 0.63<br>(0.59-0.67) | LR | 0.68<br>(0.6-0.76) | 0.66<br>(0.56-0.76) | 0.65<br>(0.61-0.69) |
| Ceftazidime/avibactam | **MLP** | **0.86**<br>(0.74-0.98) | **0.7**<br>(0.58-0.82) | **0.36**<br>(0.18-0.54) | **RF** | **0.86**<br>(0.74-0.98) | **0.71**<br>(0.55-0.87) | **0.33**<br>(0.19-0.47) | **SVM** | **0.86**<br>(0.76-0.96) | **0.72**<br>(0.58-0.86) | **0.34**<br>(0.2-0.48) |
| Ceftolozane/tazobactam | **MLP** | **0.87**<br>(0.79-0.95) | **0.69**<br>(0.57-0.81) | **0.43**<br>(0.23-0.63) | SVM | 0.83<br>(0.71-0.95) | 0.67<br>(0.53-0.81) | 0.41<br>(0.31-0.51) | SVM | 0.83<br>(0.71-0.95) | 0.67<br>(0.53-0.81) | 0.42<br>(0.3-0.54) |

To understand the contribution of specific features generated during dynamic binning, we utilized the Shapley Additive Explanations (SHAP) algorithm to determine the 50 most important features for each model [19]. As expected, features within narrow bin widths (1-20 Da) made up the greatest proportion of the 50 most important features, ranging from 30% in colistin to 76% in piperacillin/tazobactam (Supplementary Fig. 1). However, features within wider bin widths were also observed to contribute to model performance. Notably, bin widths in the range of 81-100 Da were present in the top 50 most important features for 7 out of 11 models. This supports the inclusion of signals in low intensity regions in the feature set.

### Machine learning approach using dynamic binning achieves high performance in an external dataset

We validated our approach using the DRIAMS dataset, an external MALDI-TOF MS dataset with 4139 *P. aeruginosa* spectra with corresponding AST data [17]. In contrast to our dataset, AST was performed using various methods (VITEK 2, bioMérieux; MIC Test Strips, Liofilchem; disc diffusion, ThermoFisher), the susceptibility patterns were different (Supplementary Table 1) and average spectral intensity values were lower (Supplementary Fig. 2).

With these caveats in mind, we used dynamic binning and the same methodology described above to train ML models for the 8 antimicrobials with AST labels from both datasets. We used differing combinations of Alfred and DRIAMS data in training/testing datasets (Fig. 4) with several important findings. Dynamic binning also demonstrated efficacy for the DRIAMS dataset, achieving AUROC of 0.65-0.88. Model performance varied according to which datasets were used for training and testing ranging from AUROC 0.47 to 0.88. Training the model with one dataset then testing on a different dataset adversely affected performance. Conversely, incorporating maximal data (i.e. both datasets) for model training improved performance.

**Fig. 4.**
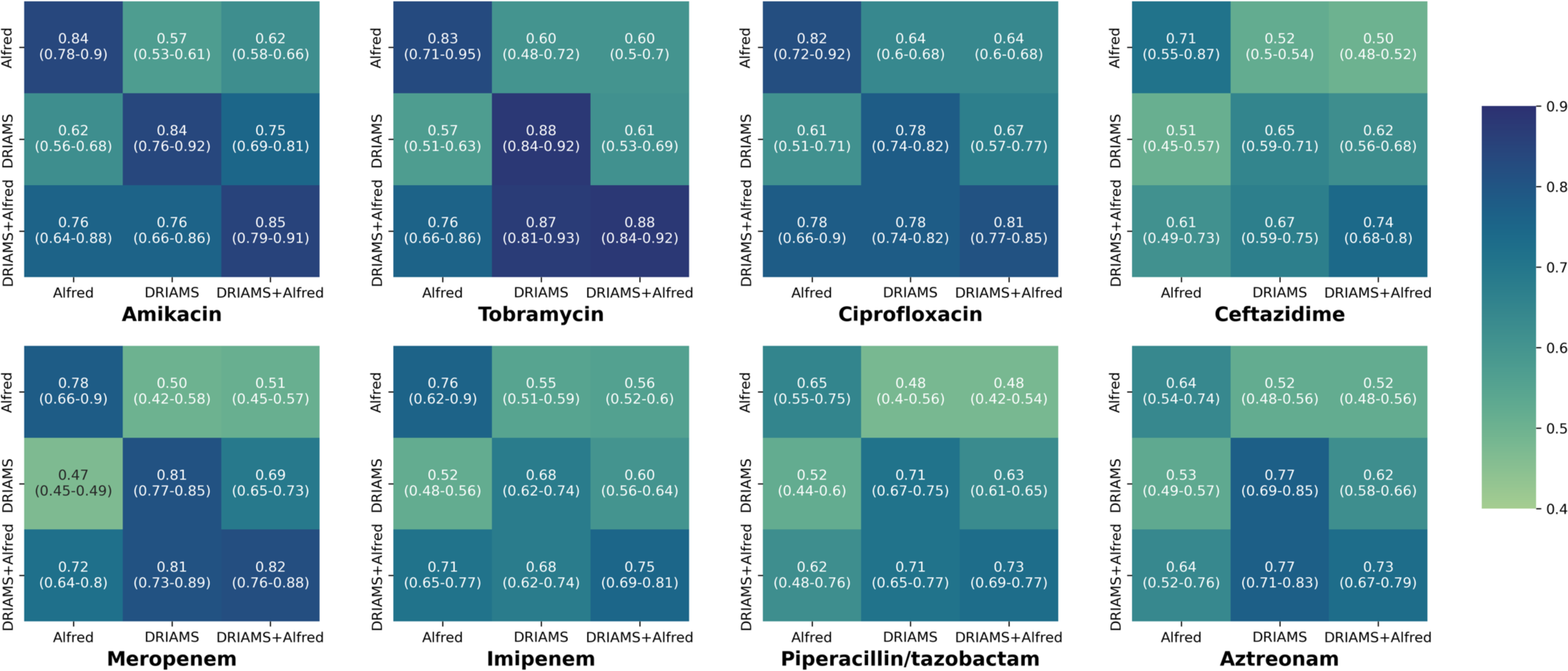
- Cross-institution performance of dynamic binning. Different training (y-axis) and testing (x-axis) sets using DRIAMS and Alfred Hospital data were used to train and evaluate the models. Data are reported as mean AUROC (95% CI) across 10 random splits. Abbreviations: AUROC – area under the receiver-operating characteristic curve; DRIAMS - Database of Resistance Information on Antimicrobials and MALDI-TOF Mass Spectra

### Further improvement in MALDI-TOF MS predictions through transfer learning

Transfer learning has been widely adopted in multiple fields of machine learning, where knowledge from one domain or task can be used to improve performance in a related domain or task [20]. In brief, rather than training a model *de novo*, a pre-trained model trained on a large dataset is used as a starting point. This pre-trained model is then fine-tuned or used as an embedding layer for a new task using a specific dataset relevant to the new task.

In order to retain key information after dimensionality reduction with dynamic binning, we used transfer learning to retrieve a compressed embedding of spectral data and used this to engineer additional features. Specifically, a vision transformer was used to develop a classification model for bacterial species (incorporating *>*20,000 spectra) (see Methods) [21]. The model achieved a testing accuracy of 0.98 (Supplementary Fig. 3).

We then applied this embedding to the *P. aeruginosa* spectra, generating an additional 256 features for each sample. We then used a stacking method to combine the learning outcomes from the dynamic binning features and the features obtained from the latent representation (see Methods). When compared to models using only dynamic binning, integrating transfer learning gave a modest but notable improvement with either equal or superior AUROC and F1 performance in 6/10 and 7/10 antimicrobials, respectively (Table 2).

**Table 2.** - Summary of model performance using transfer learning. Dynamic binning features and additional features retrieved from the latent space representation were used to train stacking models across 10 random splits. The models consisted of multiple base estimators (logistic regression, random forest, support vector machine, XGBoost, multilayer perceptron), each learned from a single data modality, and a final estimator (logistic regression) synthesizing the learning outcomes to output the final prediction. The best performances of the dynamic binning approach were used as benchmarking references. Bold text indicates when integrating transfer learning outperformed dynamic binning alone. Abbreviations: AUROC – area under the receiver-operating characteristic curve; CI – Confidence interval.

| Antimicrobial | Dynamic binning |  | Dynamic binning + Latent representation |  |
| --- | --- | --- | --- | --- |
|  | AUROC (95% CI) | F1 (95% CI) | AUROC (95% CI) | F1 (95% CI) |
| Amikacin | 0.84 (0.78-0.9) | 0.71 (0.57-0.85) | **0.86 (0.76-0.96)** | **0.72 (0.52-0.92)** |
| Tobramycin | 0.82 (0.7-0.94) | 0.7 (0.5-0.9) | **0.82 (0.74-0.9)** | 0.68 (0.54-0.82) |
| Ciprofloxacin | 0.82 (0.72-0.92) | 0.82 (0.72-0.92) | **0.82 (0.74-0.9)** | **0.83 (0.75-0.91)** |
| Aztreonam | 0.69 (0.59-0.79) | 0.52 (0.4-0.64) | 0.67 (0.57-0.77) | 0.51 (0.37-0.65) |
| Imipenem | 0.77 (0.65-0.89) | 0.74 (0.64-0.84) | 0.74 (0.62-0.86) | 0.73 (0.61-0.85) |
| Meropenem | 0.79 (0.65-0.93) | 0.61 (0.43-0.79) | **0.8 (0.68-0.92)** | **0.64 (0.48-0.8)** |
| Piperacillin/tazobactam | 0.68 (0.58-0.78) | 0.64 (0.54-0.74) | **0.7 (0.6-0.8)** | **0.65 (0.55-0.75)** |
| Ceftazidime | 0.7 (0.64-0.76) | 0.66 (0.58-0.74) | **0.71 (0.63-0.79)** | **0.66 (0.54-0.78)** |
| Ceftazidime/avibactam | 0.86 (0.74-0.98) | 0.7 (0.58-0.82) | 0.84 (0.72-0.96) | **0.73 (0.61-0.85)** |
| Ceftolozane/tazobactam | 0.87 (0.79-0.95) | 0.69 (0.57-0.81) | 0.84 (0.74-0.94) | **0.71 (0.55-0.87)** |

### Identifying potential resistome markers with dynamic binning

Interpretability of ML models is a key aspect of facilitating their future implementation in healthcare applications. As discussed above, we used the SHAP algorithm to measure the contribution of each feature to the model’s performance then calculated the average contribution over 10 random training- testing splits. After identifying the salient features, we examined whether there were differences in the regions of interest between the resistant and nonresistant groups. In order to identify potential AMR determinants, we then queried the UniProt database for proteins corresponding to the identified features (see Methods). A limitation of the UniProt database is that there are significantly fewer ‘reviewed’ (manually curated) than ‘unreviewed’ proteins [22].

Across all antimicrobials, regions with a mass less than 10,000 Da were most frequently noted (Supplementary Table 2). Notably, 7,565-7,630 Da was the most important feature in 6 antimicrobials, 5 of which are *β*-lactams or *β*-lactam/*β*-lactamase inhibitor combinations. To further analyze differences between resistant/non-resistant groups in the 7,565-7,630 Da range, we determined highest peaks for each isolate. In the ceftazidime/avibactam analysis, resistant isolates were unimodal with 71.23% having a peak at 7,600-7,620 Da (Fig. 5). In contrast, non-resistant isolates were bimodal with peaks at 7,600-7,620 Da (39.01%) and 7,570-7,590 Da (19.87%), leading to significantly lower mean peak intensity (*p*=1.6×10^-5^) in the 7,600-7,620 Da region. These findings were also observed for meropenem and ceftolozane/tazobactam *(p*=6.3×10^-5^ and 4.6×10^-4^, respectively). By querying the UniProt database of reviewed proteins, we identified protein RegB in the 7570-7590 Da region, and major cold shock protein CspA and UPF0337 protein PA4738 in the 76007620 Da region. Additionally, there were 97 ‘unreviewed’ proteins in those regions. The complete list can be found in Supplementary File 2.

**Fig. 5.**
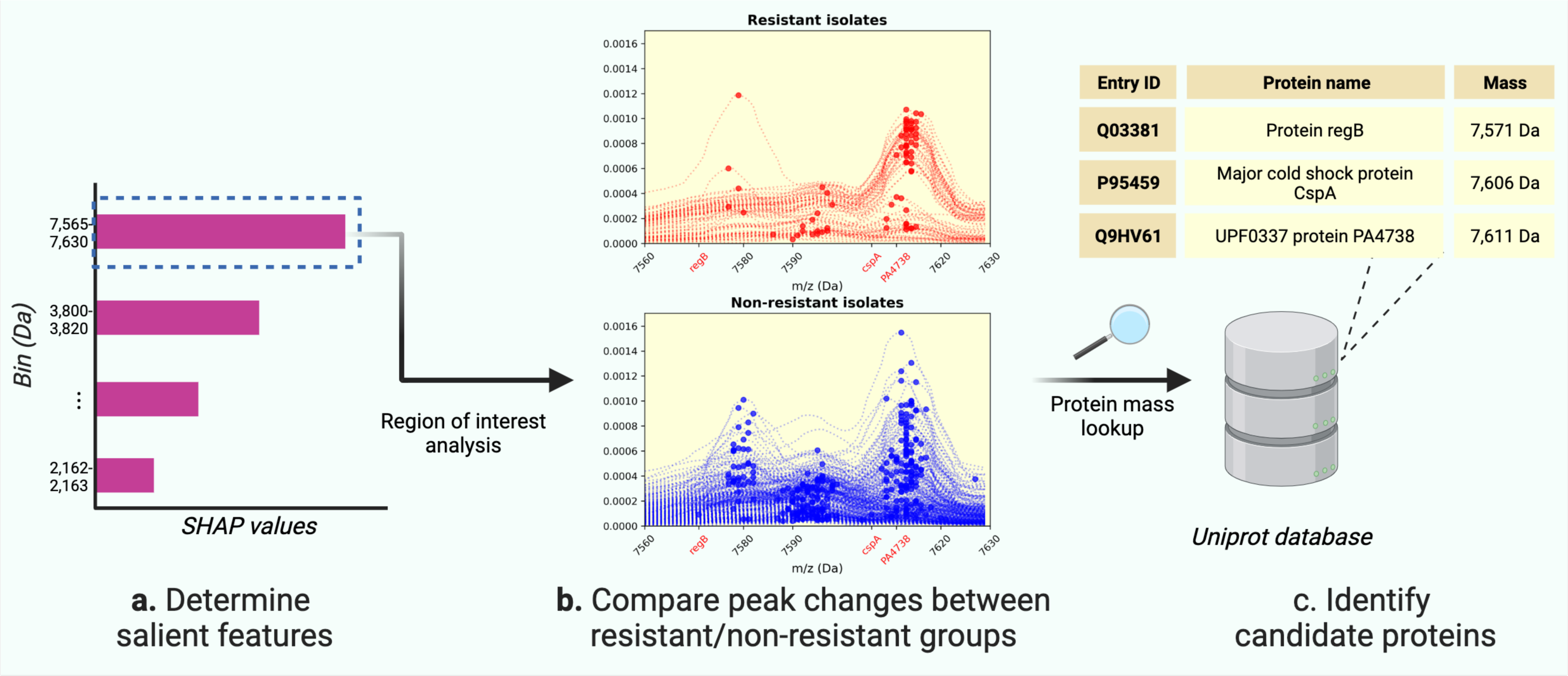
- Case study of interpretability for models predicting ceftazidime/avibactam resistance. a. Salient features were identified with SHAP algorithm b. Each red/blue line illustrates intensity values in the regions of interest for one isolate. The dots indicate the highest peak within the region. Red text (x-axis) indicates the theoretical mass (Da) of the mapped proteins in step c. Here, we only display results for the ‘reviewed’ proteins. Spectral comparison for other antimicrobials can be found in Supplementary Fig. 4. Abbreviations: m/z – mass-to-charge ratio; Da – Dalton; SHAP - Shapley Additive Explanation

Protein RegB facilitates the production of exotoxin A, one of the most powerful virulence factors of *P. aeruginosa* [23]. With protein CspA, there is correlation between its presence and the survival rate in extreme weather environments [24]. The PA4738 gene encodes a hydrophilin protein, which is expressed under conditions of osmotic stress and may be associated with antibiotic tolerance in *P. aeruginosa* [25, 26]. Therefore, PA4378 may be a potential resistance determinant for these *β*-lactam antimicrobials.

## Discussion

In this study, we successfully used novel machine learning methods to predict AMR for *P. aeruginosa* from MALDI-TOF MS data. Despite the complexity of resistance mechanisms in *P. aeruginosa*, we achieved high predictive performance using five different ML approaches across 11 antimicrobials, including novel *β*-lactam/*β*-lactamase inhibitor agents. We also showed validity of our approach with comparable high-performance using *P. aeruginosa* MALDI-TOF MS data from a publicly available dataset. While several previous studies have utilized MALDI-TOF MS for AMR prediction in other bacteria, this has not been done for *P. aeruginosa*. MALDI-TOF MS data are now routinely generated in clinical microbiology for bacterial identification testing and our work demonstrates that these data could be successfully re-purposed for rapidly inferring AST without additional costs.

The performance of our models varied depending on the antimicrobial type, with AUROC ranging from 0.55 to 0.92, consistent with previous studies using MALDI-TOF MS to predict AMR [15, 17]. Our models performed well for fluoroquinolones and aminoglycosides (AUROC 0.82-0.84), which aligns with previous findings, using not only MALDI-TOF MS but also genomic data [15, 27–30]. This may reflect the more limited number of AMR mechanisms in fluoroquinolones and aminoglycosides [31, 32]. In contrast, *β*-lactams showed more variable performance (AUROC 0.68-0.87), reflecting the complexity of AMR mechanisms. This has also been noted in *K. pneumoniae*, with AUROC for carbapenems ranging from 0.55 to 0.95. The heterogeneity of these findings highlights the challenge in developing a reliable AMR prediction model using MALDI-TOF MS.

In comparison to a previous genomic study [29], we achieved superior AUROC for amikacin (0.84 vs 0.79) and comparable results for meropenem (0.79 vs 0.79). This is highly encouraging, considering the lower cost and labor associated with obtaining MALDI-TOF spectra compared to genomic data [33]. Integrating multiple data modalities has the potential to improve performance and highlights a potential future direction for enhancing AMR prediction models. For instance, Khaledi et al. [27] effectively combined gene expression information with genomics, achieving F1-macro scores of 0.82-0.92.

The differing performance of MALDI-TOF MS AMR predictions across antimicrobial classes noted in our study has distinct clinical implications. MALDI-TOF MS models could be used as a rule-out test for ciprofloxacin resistance due to a high recall score. This may help prevent failure in outpatient treatment given that ciprofloxacin is one of the few oral treatments for *P. aeruginosa* infection. Conversely, high precision of MALDI-TOF models may help confirm susceptibility and avoid inappropriate empirical use of last-line treatments (e.g., ceftolozane/tazobactam and ceftazidime/avibactam) in patients with resistant isolates.

Our benchmarking analysis demonstrated that dynamic binning and transfer learning improve performance. These approaches complement each other: dynamic binning reduces irrelevant features, while transfer learning adds potentially useful ones. Healthcare-related data often require feature reduction due to their high dimensionality and limited sample size [34]. However, using a wide bin width in fixed-length binning may inadvertently lead to the loss of informative intensity peaks when multiple peaks fall within the same bin. This likely explains why 20-Da binning performed less effectively than 3-Da binning. In terms of transfer learning, although there were modest improvements, models can benefit from incorporating informative features. While our pretrained model showed exceptional performance in bacterial identification, which is the primary use case of MALDI-TOF MS in routine laboratory settings, the model may perceive this task as relatively straightforward. Therefore, defining different pre-training tasks could enable the model to uncover a more beneficial latent representation. As noted in other fields, with the increasing availability of public data and state-of-the-art pre-training approaches (e.g., the GPT large language model [35]), we anticipate that transfer learning will play a significant role in developing AMR prediction models using MALDI-TOF MS data.

We demonstrated the effectiveness of dynamic binning by validating it on a large public dataset. Our models generally performed better when the training and testing sets were from the same site, consistent with Weis et al.’s findings [17]. Interestingly, Yu et al. noted that their model, which predicts methicillin-resistant *Staphylococcus aureus*, performed better on prospective data from their center than datasets from other healthcare centers [13]. This may be explained by underlying biological differences in geographically different datasets [36], such as clonal relatedness and differences in AST profiles. Furthermore, there may be variability in MALDI-TOF MS data collected from different sources due to multiple technical factors such as sample preparation or device calibration [37, 38]. While including additional training data can improve model performance, incorporating an external dataset may introduce noise and degrade model performance. Therefore, to ensure the utility of ML models, it is crucial to reevaluate the training process, considering the dynamic changes in pathogen characteristics over time.

Interpreting ML models is challenging due to their complexity and performance discrepancies across datasets. In this study, we used the SHAP algorithm for post-hoc interpretability [19]. Our findings indicate that the presence or absence of peaks significantly influences the model’s performance, consistent with previous work [13, 17, 39]. Interpreting these findings is challenging for several reasons. Firstly, there are limited data on changes in MALDI-TOF spectra related to AMR in *P. aeruginosa*. Additionally, the large numbers of uncharacterized proteins make it difficult to explain key features. Literature-based manual curation requires substantial time and effort, contrasting with the rapid generation of numerous newly identified protein entities through high-throughput methods [22]. Finally, the current m/z (mass-to-charge ratio) range (2,000-20,000 Da) may be inadequate for studying well-characterized AMR genes in *P. aeruginosa*, as most resistance determinants have higher masses. For example, the ampC *β*-lactamase (UniProt ID: P24735) and oprD (ID: P32722), a membrane porin involved in carbapenem uptake, have masses of 43,401 Da and 48,360 Da, respectively. This limitation extends beyond *P. aeruginosa* to other species [40–42].

We acknowledge several limitations in our study. Firstly, our dataset is relatively small, making it challenging to apply cutting-edge deep learning architectures that typically require a larger training dataset. Secondly, putative resistance determinants identified through model interpretation should be validated experimentally [43]. In our study, multiple proteins had similar masses, making it difficult to identify exact AMR biomarkers. This problem could be overcome by molecular docking simulations to analyze changes in antimicrobial binding affinity [13] or incorporating genomic data [44]. Lastly, further prospective evaluation is crucial prior to implementing predictive models in a clinical setting. To date this has been limited to retrospective studies that evaluate potential deleterious outcomes where there are disagreements between clinical decisions and the predicted outcomes of ML models [17, 39].

In conclusion, our work provides the first insight into applying AI approaches to predict AMR in *P. aeruginosa* using MALDI-TOF MS data. Our findings show that model improvement is achieved through efficient feature engineering and leveraging extensive datasets. Future directions include evaluating dynamic binning with high-risk pathogens and integrating multiple data modalities, not only to improve predictive performance, but also foster explainable AI.

## Methods

### MALDI-TOF MS data collection

The study was reviewed and approved by the Alfred Hospital Ethics Committee. We included all clinical *P. aeruginosa* isolates from an institutional collection at the Alfred Hospital, Melbourne, Australia. The Alfred Hospital is a 638-bed quaternary hospital, with a cystic fibrosis and lung transplant state referral service. Isolates were collected from 2004-2021. Mass spectra were obtained using Microflex Biotyper^®^ (Bruker Daltonics). Only *P. aeruginosa* isolates with an identification log score equal or greater than 2 were included.

### Antimicrobial susceptibility testing

We conducted AST for all isolates by performing broth microdilution using the Sensititre^TM^ (ThermoFisher) Gram negative plate (plate code: DKMGN). The reported MIC values were then converted to binary labels using the European Committee on Antimicrobial Susceptibility Testing version 13 (EUCAST v13.0) breakpoint tables [16]. Specifically, we classified isolate as resistant (1) if the MIC value is greater than (*>*) the resistant breakpoint, and as non-resistant (0) if the MIC value is less than or equal to (≤) the resistant breakpoint.

### Spectra preprocessing

We followed the 5-step workflow outlined in [17]. In brief, the pipeline includes: (1) variance stabilization with square-root transformation; (2) data smoothing with the Savitzky-Golay filter; (3) baseline removal with SNIP; (4) Total-Ion-Current (TIC) calibration; (5) and 2000-20000 Da range trimming. Preprocessed spectra will be either binned with dynamic binning or fixed-length (3-Da and 20-Da) binning (Fig. 2).

### Dynamic binning method

Firstly, we computed the average peak counts for 500-Da disjoint sub-regions. With the preprocessed spectra, we used MALDIquant to detect intensity peaks [45]. For each sample, we counted the number of peaks appearing in each subregion, and finally calculated the average number of peaks across the whole dataset (Supplementary Fig. 5). Secondly, we determined a bin width value, ranging from 1 to 100 Da, for each sub-region, which is inversely proportional to its corresponding average peak count.

In contrast to the binning process in [17], which used mean values, we computed the maximum value of each bin. This was done as bin widths in dynamic binning can be much larger than in fixed-length binning, making it possible for a high intensity value to be diluted out by a large number of low intensity values if a mean is used. Following both dynamic binning and fixed-length binning, we obtained a tabular data representation where each row represents a sample and the columns represent the maximum intensity of each bin.

### Machine learning model development

Firstly, the dataset was randomly split into a training (80%) and testing (20%) set. We then used Scikit-learn and its related packages to build our models [46, 47]. We used the following machine learning approaches: LR, RF, SVM, XGB, and MLP with hyperparameter tuning to select the best-performing hyperparameter set for each model. Specifically, we tuned the hyperparameters with 5-fold cross validation on the training set in order to maximize AUROC score. For each tunable hyperparameter, we used Optuna to determine the optimized value after 100 trials. Based on the history of the trial records, Optuna used a Bayesian approach, namely Tree-structured Parzen Estimator, to decide the next value to assess [47]. Details of hyperparameters can be found in our code repository (train.py file). For performance evaluation, we predicted AMR for the testing set using the fine-tuned models. Performance metrics included accuracy, AUROC, F1-score, precision, recall, and log-loss [48]. In addition, we reported results of true negative (true susceptible), false negative (false susceptible), true positive (true resistant), and false positive (false resistant).

### External dataset assessment

We identified 4139 *P. aeruginosa* isolates in the publicly available DRIAMS dataset [17]. These isolates had both MALDI-TOF MS spectra and AST data for 8 antimicrobials: amikacin, tobramycin, ciprofloxacin, ceftazidime, meropenem, imipenem, piperacillin-tazobactam, and aztreonam. We followed the same process as described above to preprocess the raw spectra, extract features, train and evaluate the models. For each antimicrobial we assessed 3 permutations of using different datasets (only Alfred Hospital data, only DRIAMS data, and all data) for training and testing, resulting in 9 different settings for each antimicrobial.

### Bacterial identification with vision transformer

We built a vision transformer model to classify high-risk pathogens, including *Enterobacter cloacae*, *Staphylococcus aureus*, *Klebsiella pneumoniae*, *Acinetobacter baumannii*, *Pseudomonas aeruginosa*, and *Enterococcus faecium*. Each spectrum was viewed as a 1-D image. We used the 1-Da binning features to enhance resolution. Thus, the input shape (channel, height, width) is (1, 1, 18000).

The vision transformer model consisted of 8 attention heads, and the final hidden layer has 256 units. We used cross entropy loss as the loss function, and the models were trained for 1000 epochs with a batch size of 128, learning rate of 10^-3^, and dropout rate of 0.2. The model from the best checkpoint, determined via validation loss, was selected as the final model. We used Pytorch and Pytorch-lightning for the training and testing process [49, 50]. For latent representation visualization, we used t-SNE to project the embedding into two-dimensional space [51].

### AMR prediction with stacking method

With stacking ensemble learning, the predicted labels generated by multiple base estimators are used as features for the final estimator (Supplementary Fig. 6). Here, we utilized Scikit-learn’s StackingClassifier to develop the stacking model, with LR, RF, SVM, XGB, and MLP as the base estimators. Instead of learning from the whole combined feature set, for each ML approach, we created 2 distinct base models, one learns from the dynamic binning features, and the other learns from the latent embedding features, resulting in a total of 10 different features for each sample. For the final estimator, we used the default LR model.

### Model interpretation with SHAP algorithms

To calculate the feature importance score in dynamic binning, we ran SHAP algorithm for each run of the best-performing model per antimicrobial. Briefly, the algorithm computes the contributing score of each individual feature by measuring the difference of model’s performance when the corresponding feature is included or excluded in different permutations of feature sets. We used TreeExplainer for XGB and RF, and KernelExplainer for LR, SVM and MLP [19, 52]. We then calculated the average SHAP values from 10 random splits to identify the most important features for the best classifiers per antimicrobial. To identify protein candidates, we filtered the *P. aeruginosa* proteins available in the UniProt database by checking whether their theoretical masses are within the bin of interest. The list of *P. aeruginosa* proteins and their corresponding masses were obtained using UniProt’s search and download functions.

## Data availability

MALDI-TOF spectral data and broth microdilution results are available at Figshare. All samples were de-identified.

## Code availability

The source code is available on GitHub.

## Acknowledgments

The authors thank B. Vezina (Department of Infectious Diseases, Monash University/Alfred Hospital) for discussion around developing dynamic binning.

## Author contributions

HA.N, N.M. and A.Y.P. conceived the study. J.A.W. designed and supervised sampling and collection of bacterial isolates. J.A.W., L.V.B., G.Z.B, H.Z. and R.T. collected the bacterial isolates, performed bacterial characterization and generated MALDI-TOF spectra. HA.N. and B.A. preprocessed MALDI-TOF spectra. HA.N., J.S., D.L.D., G.I.W. and N.M. conceptualized machine learning analyses. HA.N. developed machine learning models and evaluated predictive performances. HA.N. and N.M. analyzed all results. HA.N. and N.M. wrote the manuscript with comments and feedback of all of the co-authors. All authors read and approved the manuscript.

