## Supplementary Information for "Predicting *Pseudomonas aeruginosa* drug resistance using artificial intelligence and clinical MALDI-TOF mass spectra"

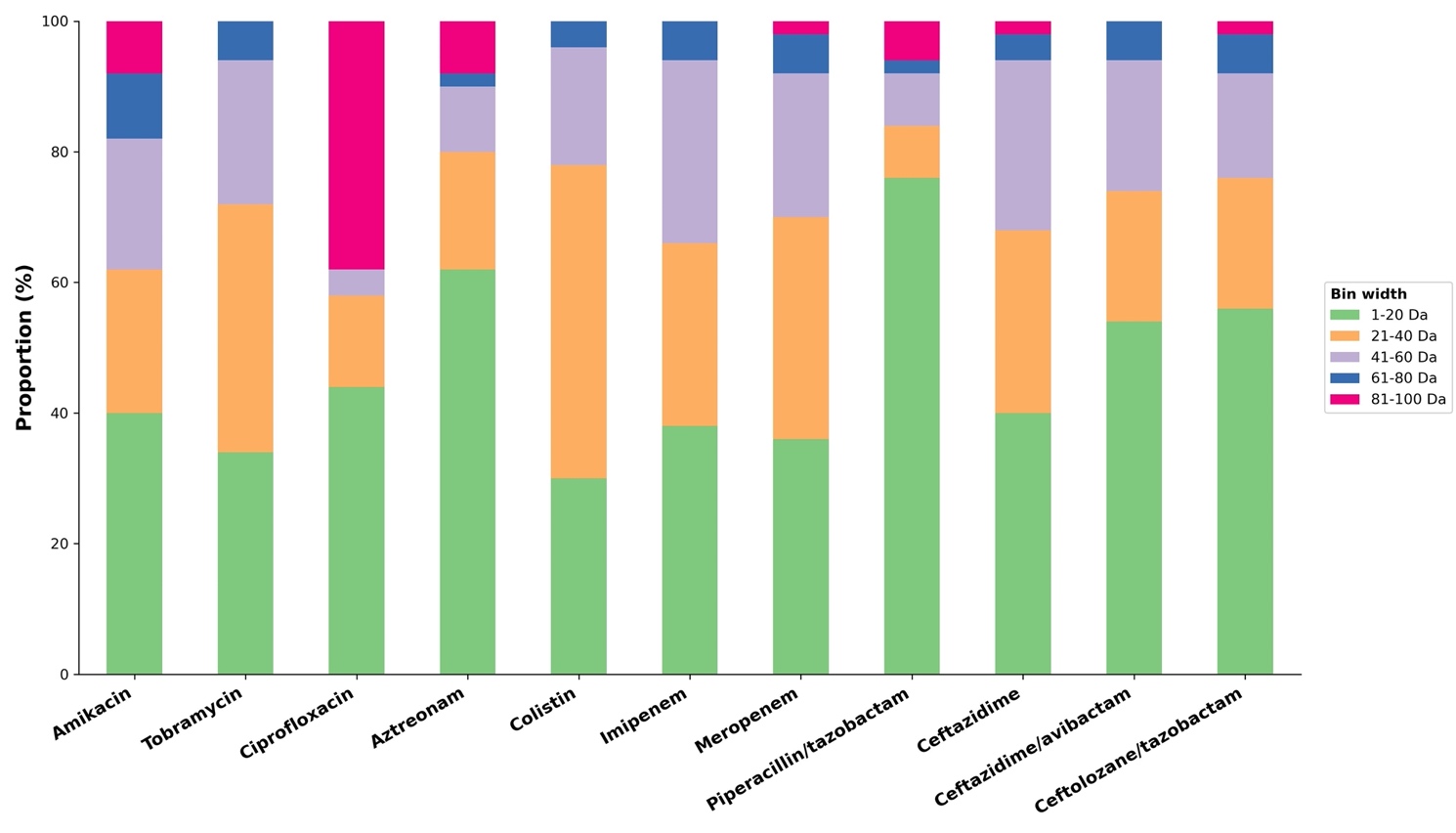

**Supplementary Fig. 1 - Distribution of bin width of salient features.** Bin widths of the top 50 most important features are grouped into contiguous 20-Da bin.

Abbreviations: Da – Dalton.

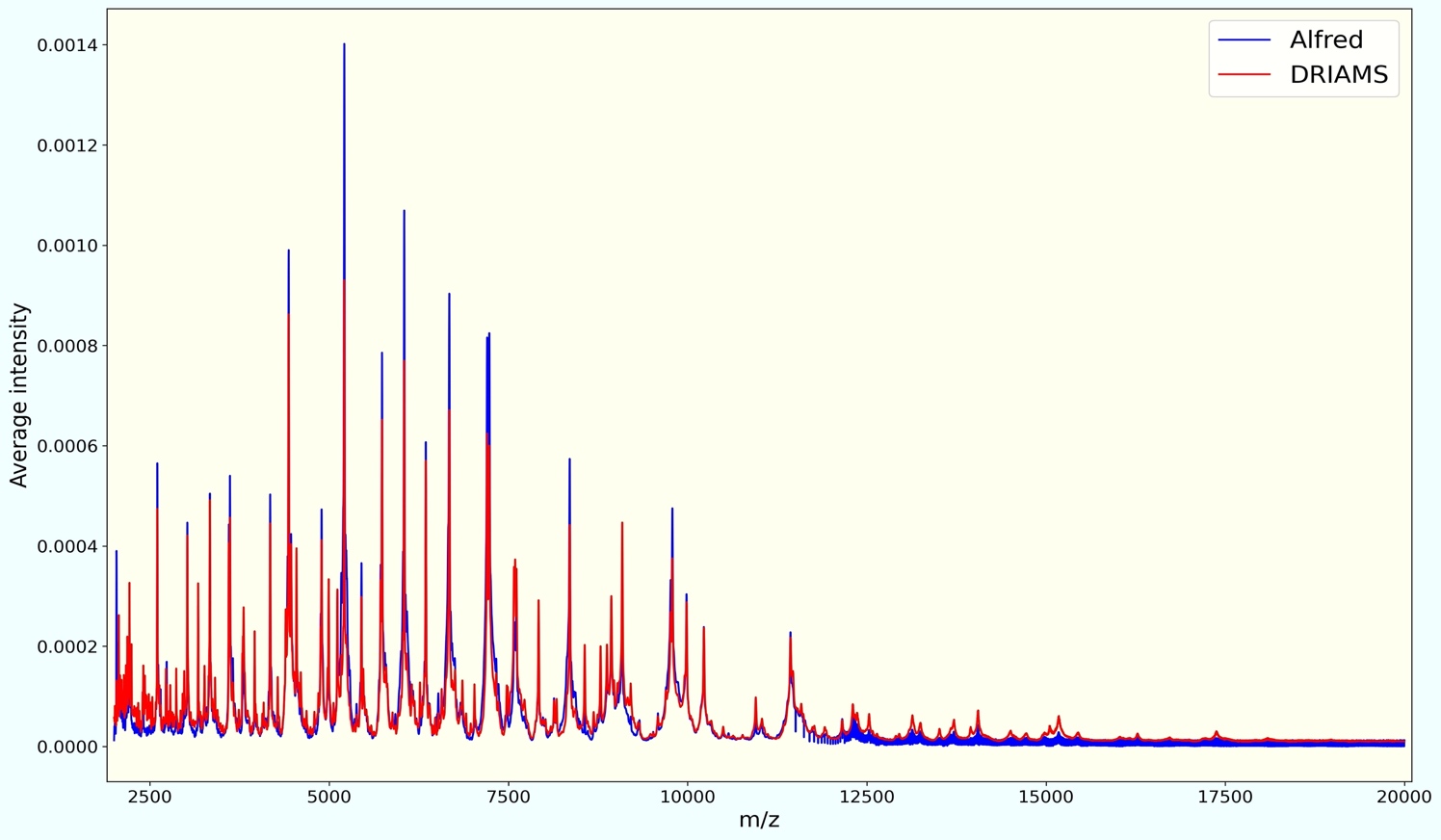

**Supplementary Fig. 2 - Differences in peak intensity between DRIAMS and Alfred Hospital datasets.** Mean intensity values of MALDI-TOF spectra of *P. aeruginosa* from the DRIAMS and Alfred Hospital datasets are shown. All MALDI-TOF spectra were preprocessed and trimmed to 2,000-20,000 Da. We used 1-Da binning to generate a uniformed intensity vector of 18,000 dimensions for each isolate.

Abbreviations: DRIAMS - Database of Resistance Information on Antimicrobials and MALDI-TOF Mass Spectra; m/z – mass-to-charge ratio.

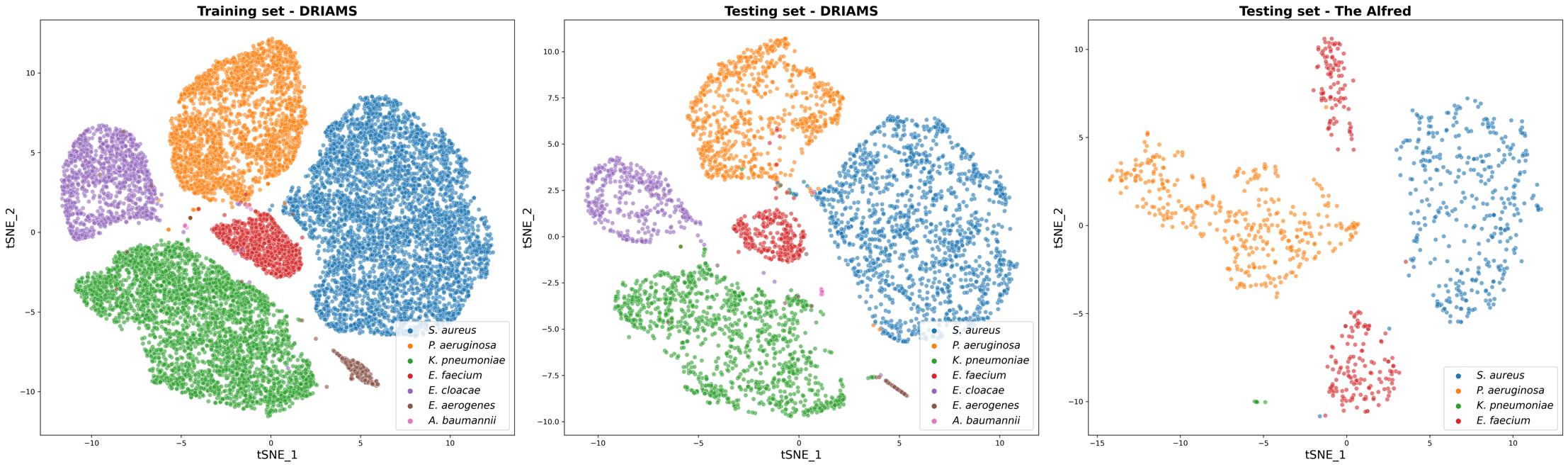

**Supplementary Fig. 3 - Visualization of latent representation.** We used t-distributed Stochastic Neighbor Embedding (t-SNE) to project the hidden layer retrieved from the vision transformer model into 2-D space. Shown are the clusters of pathogens in the training set (left, DRIAMS dataset), testing set (center, DRIAMS dataset), and Alfred Hospital dataset (right).

Abbreviations: DRIAMS - Database of Resistance Information on Antimicrobials and MALDI-TOF Mass Spectra; tSNE - t-distributed Stochastic Neighbor Embedding; *S. aureus* – *Staphylococcus aureus*; *P. aeruginosa* – *Pseudomonas aeruginosa*; *K. pneumoniae* – *Klebsiella pneumoniae*; *E. faecium* – *Enterococcus faecium*; *E. cloacae* – *Enterobacter cloacae*; *E. aerogenes* - *Enterobacter aerogenes*; *A. baumannii* – *Acinetobacter baumannii.*

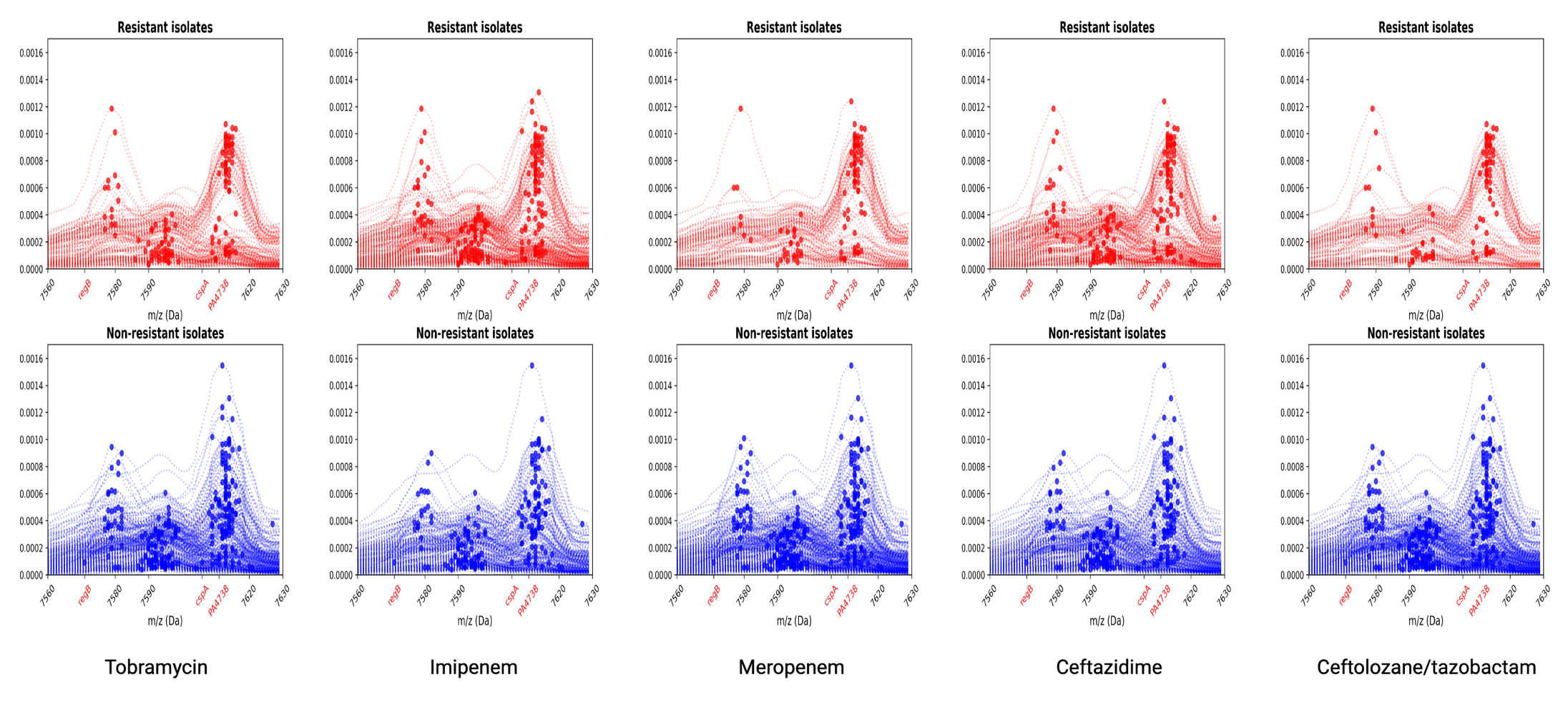

**Supplementary Fig. 4 - Spectral comparison for other antimicrobials.** In all presenting antimicrobials, the feature bin of 7565-7630 Da is the highest contributing feature for the AMR prediction models.

Abbreviations: m/z – mass-to-charge ratio; Da – Dalton.

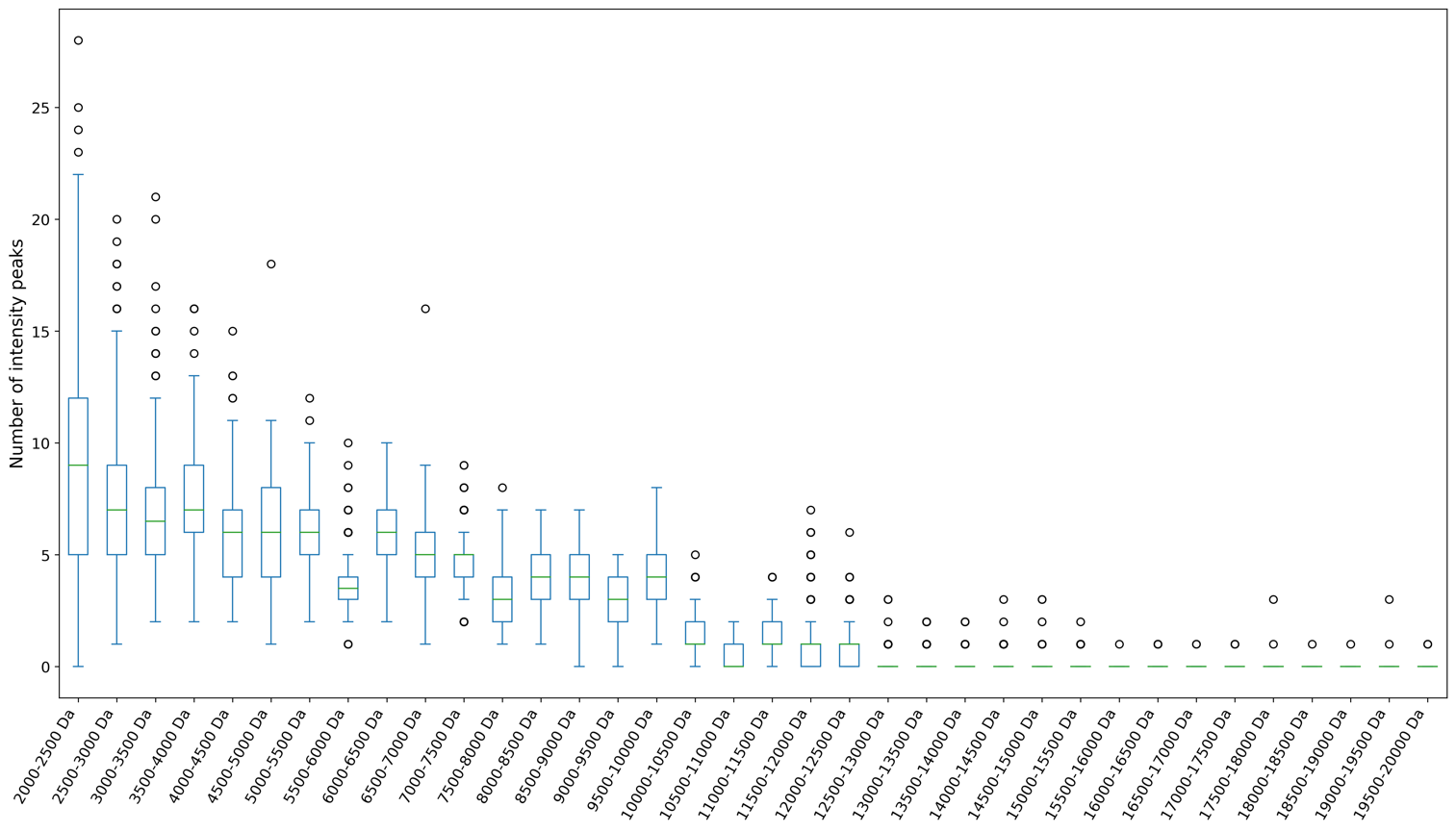

**Supplementary Fig. 5 - The number of intensity peaks per 500-Da region.** Boxplots illustrate distribution of peak count across all spectra.

Abbreviations**:** m/z – mass-to-charge ratio.

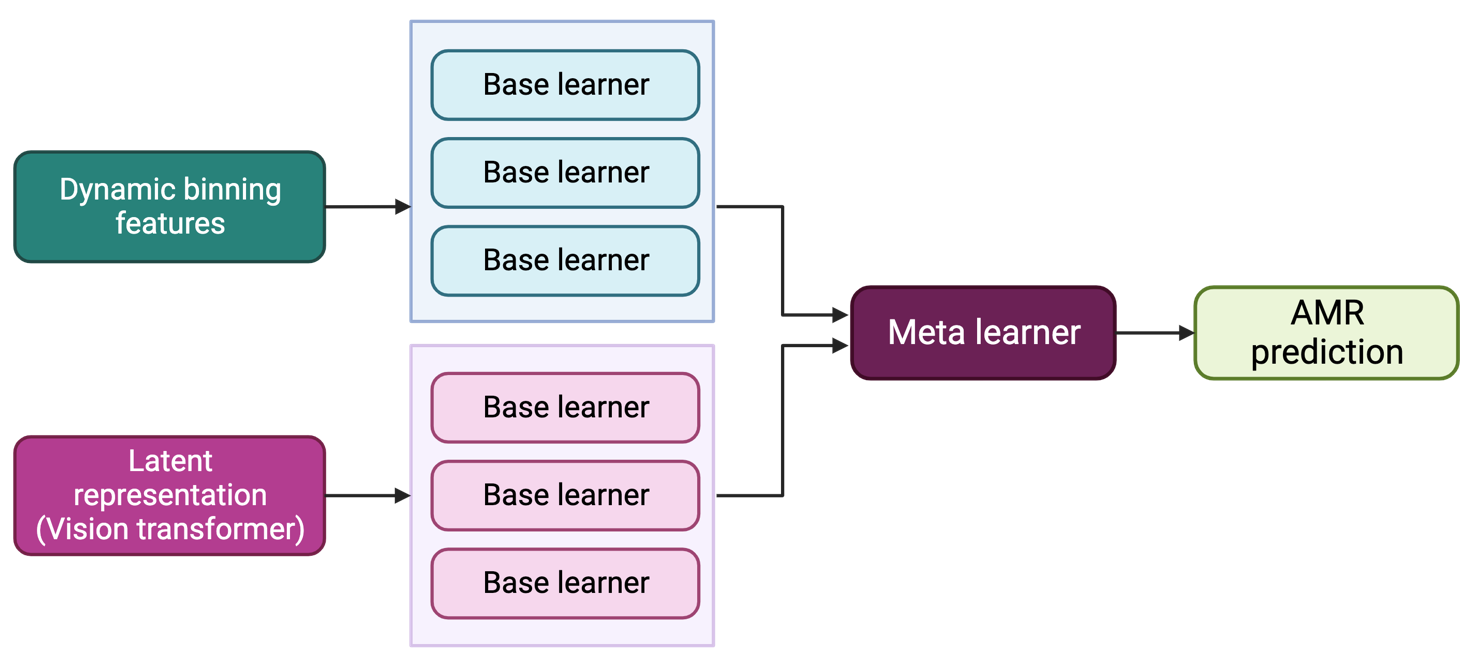

**Supplementary Fig. 6 - High-level architecture of stacking model.** Each input data type, i.e., dynamic binning features and latent representation from vision transformer model, was trained separately with different base learners.

Abbreviations: AMR – antimicrobial resistance.

**Supplementary Table 1 - AST phenotypes of *P. aeruginosa* isolates in DRIAMS dataset.** Data were collected from multiple healthcare centers using different antimicrobial susceptibility testing assay methods. This resulted in varying numbers of isolates tested for each antimicrobial.

| **Antimicrobial** | **Sample count** | **Resistant** | **Non-resistant** |
| --- | --- | --- | --- |
| Amikacin | 3141 | 106 (3.37%) | 3035 (96.63%) |
| Tobramycin | 3489 | 258 (7.39%) | 3231 (92.61%) |
| Ciprofloxacin | 3962 | 740 (18.68%) | 3222 (81.32%) |
| Ceftazidime | 3281 | 293 (8.93%) | 2988 (91.07%) |
| Meropenem | 3554 | 336 (9.45%) | 3218 (90.55%) |
| Imipenem | 3170 | 420 (13.25%) | 2750 (86.75%) |
| Piperacillin/tazobactam | 3965 | 676 (17.05%) | 3289 (82.95%) |
| Aztreonam | 987 | 212 (21.48%) | 775 (78.52%) |

**Supplementary Table 2 - Salient features for predictive performance.** For each antimicrobial, we report 10 features achieving the highest average SHAP values across 10 running times. Ordinal numbers indicate the corresponding SHAP value ranks.

| **Antimicrobial** | **1^th^** | **2^nd^** | **3^rd^** | **4^th^** | **5^th^** | **6^th^** | **7^th^** | **8^th^** | **9^th^** | **10^th^** |
| --- | --- | --- | --- | --- | --- | --- | --- | --- | --- | --- |
| Amikacin | 5500-5560 | 6408-6442 | 5434-5465 | 5372-5403 | 2724-2740 | 6374-6408 | 2429-2430 | 2430-2431 | 7343-7392 | 4408-4442 |
| Tobramycin | 7565-7630 | 5680-5740 | 5434-5465 | 5740-5800 | 4533-4566 | 9069-9138 | 4408-4442 | 6680-6725 | 3288-3312 | 6340-6374 |
| Ciprofloxacin | 2724-2740 | 3720-3740 | 4408-4442 | 2628-2644 | 2548-2564 | 5500-5560 | 5341-5372 | 11590-11680 | 19800-19900 | 19600-19700 |
| Aztreonam | 6476-6500 | 2458-2459 | 5740-5800 | 9555-9610 | 4764-4797 | 4238-4272 | 12698-12797 | 3640-3660 | 2252-2253 | 2175-2176 |
| Ceftazidime | 7565-7630 | 5186-5217 | 5680-5740 | 5434-5465 | 6680-6725 | 2033-2034 | 6034-6068 | 6340-6374 | 9775-9830 | 8300-8360 |
| Ceftazidime/avibactam | 7565-7630 | 3800-3820 | 5434-5465 | 5403-5434 | 2167-2168 | 2166-2167 | 4896-4929 | 2165-2166 | 9775-9830 | 2168-2169 |
| Ceftolozane/tazobactam | 7565-7630 | 5740-5800 | 3800-3820 | 5403-5434 | 2167-2168 | 5186-5217 | 4896-4929 | 2033-2034 | 2166-2167 | 5500-5560 |
| Piperacillin/tazobactam | 5500-5560 | 2644-2660 | 2437-2438 | 2628-2644 | 2436-2437 | 2548-2564 | 2724-2740 | 2377-2378 | 5680-5740 | 4408-4442 |
| Meropenem | 7565-7630 | 5740-5800 | 5680-5740 | 5186-5217 | 4408-4442 | 2037-2038 | 9775-9830 | 3288-3312 | 5434-5465 | 3800-3820 |
| Imipenem | 7565-7630 | 5434-5465 | 9069-9138 | 6680-6725 | 5217-5248 | 5680-5740 | 5740-5800 | 9720-9775 | 4408-4442 | 9775-9830 |
